# High-fidelity Database-free Deep Learning Reconstruction for Real-time Cine Cardiac MRI

**DOI:** 10.1101/2023.02.13.528388

**Authors:** Ömer Burak Demirel, Chi Zhang, Burhaneddin Yaman, Merve Gulle, Chetan Shenoy, Tim Leiner, Peter Kellman, Mehmet Akçakaya

## Abstract

Real-time cine cardiac MRI provides an ECG-free free-breathing alternative to clinical gold-standard ECG-gated breath-hold segmented cine MRI for evaluation of heart function. Real-time cine MRI data acquisition during free breathing snapshot imaging enables imaging of patient cohorts that cannot be imaged with segmented or breath-hold acquisitions, but requires rapid imaging to achieve sufficient spatial-temporal resolutions. However, at high acceleration rates, conventional reconstruction techniques suffer from residual aliasing and temporal blurring, including advanced methods such as compressed sensing with radial trajectories. Recently, deep learning (DL) reconstruction has emerged as a powerful tool in MRI. However, its utility for free-breathing real-time cine MRI has been limited, as database-learning of spatio-temporal correlations with varying breathing and cardiac motion patterns across subjects has been challenging. Zero-shot self-supervised physics-guided deep learning (PG-DL) reconstruction has been proposed to overcome such challenges of database training by enabling subject-specific training. In this work, we adapt zeroshot PG-DL for real-time cine MRI with a spatio-temporal regularization. We compare our method to TGRAPPA, locally low-rank (LLR) regularized reconstruction and database-trained PG-DL reconstruction, both for retrospectively and prospectively accelerated datasets. Results on highly accelerated real-time Cartesian cine MRI show that the proposed method outperforms other reconstruction methods, both visibly in terms of noise and aliasing, and quantitatively.

## I. INTRODUCTION

Real-time cine cardiac MRI is a free-breathing and ECG-free alternative to gold-standard ECG-gated and breathhold cine MRI for functional and volumetric assessment of the heart to diagnose cardiovascular diseases [1]. ECG-gated acquisitions require breath-hold imaging to minimize respiratory motion artifacts, but it may fail in patients with arrhythmias or difficulty breath-holding, and is not possible during exercise stress [2]. On the other hand, real-time cine MRI does not require ECG gating or breath-holding, which appeals to a more general patient cohort [3]. However, to achieve sufficient spatio-temporal resolutions, highly accelerated data acquisition is needed. This imposes significant challenges for subsequent image reconstruction.

Previous approaches to real-time cine MRI utilize parallel imaging in conjunction with Cartesian [4], [5] or non-Cartesian trajectories, especially radial imaging [6]. Subsequently, compressed sensing (CS) with radial sampling was used to further improve spatio-temporal resolution [7]. Yet, at very high acceleration rates, CS reconstruction may suffer from residual aliasing artifacts and blurring [8]. Recently physics-guided deep learning (PG-DL) reconstruction has emerged as a powerful method for highly-accelerated MRI, improving on parallel imaging and CS [9]–[12]. The utility of DL reconstruction in real-time cine MRI has so far been limited to data-driven image enhancement approaches that learn a mapping between aliased and artifact-free images [13], [14]. However, learning a spatio-temporally regularized reconstruction for free-breathing real-time cine MRI has its own challenges, particularly due to the varying breathing and cardiac motion patterns between subjects, along with a lack of fully-sampled reference data for highly-accelerated scans.

In this study, we propose to use subject-specific zeroshot self-supervised PG-DL [15] without a training database or ground-truth data to improve highly accelerated realtime Cartesian cine MRI. The proposed PG-DL network operates on the subject of interest using spatio-temporal regularization. The proposed approach was compared with TGRAPPA [4], locally-low-rank (LLR) regularized reconstruction [16] and database-trained self-supervised PG-DL [12] with both retrospectively and prospectively accelerated datasets. Results show that the subject-specific zero-shot PG-DL reconstruction substantially improves upon other methods, showing excellent image quality compared to baseline images while preserving temporal fidelity. Results on retrospectively highly-accelerated real-time cine MRI show that the proposed method improves on existing methods in terms of PSNR (up to 35.98% gain) and SSIM (up to 39.29% gain) when compared to standard acceleration. Additionally, our approach establishes the feasibility of real-time Cartesian cine MRI with 8-fold prospective acceleration that matches the spatio-temporal resolution of breath-hold cine MRI.

## II. METHODS

### A. Physics-Guided Deep Learning MRI Reconstruction

Regularized MRI reconstruction is formulated as:

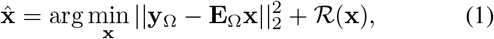

where **x** is the image of interest, **y_Ω_** is the acquired multichannel k-space with Ω is the undersampling pattern, **E_Ω_** is the multi-coil encoding operator and 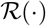 is the regularizes PG-DL methods typically solve Eq. 1 by algorithm unrolling that alternates between conventional data-fidelity (DF) with acquired k-space and a proximal operator that is implicitly implemented via neural networks. Among various techniques to decouple the DF and regularizer terms [17], proximal gradient descent alternates between:

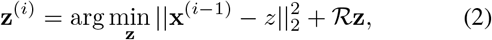

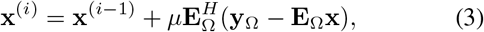

where **z** is an auxiliary variable at the the *i^th^* iteration and *μ* is penalty term. These networks are typically trained end-to-end in a supervised manner using reference images [11].

### B. Database-free Self-supervised PG-DL Reconstruction

Several unsupervised learning strategies have been proposed to tackle the challenges associated with the end-to-end supervised training of unrolled networks that require reference data [18]. Among these, self-supervised learning strategies have been utilized in various applications [12], [19]–[21]. Nonetheless, these require a database for training, which is hard to curate for real-time cine MRI due to breathing pattern differences among subjects. Recently, zeroshot self-supervised learning via data undersampling (ZS-SSDU) has been proposed to enable database-free training of PG-DL reconstructions [15]. ZS-SSDU splits the acquired k-space locations, Ω, into three disjoint sets. First two sets are similar to database-trained SSDU [12]: Ω used in the data fidelity units of the unrolled network, Λ used to define selfsupervised loss. The third set Γ defines self-validation loss to determine an early stopping criteria and avoid overfitting. Using multi-mask data-augmentation strategy [22], the ZS-SSDU training loss is given as:

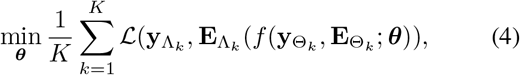

where ***θ*** denotes the network parameters, *f* (**y**_Θ_*k*__, **E**_Θ_*k*__; ***θ***) is the network output for input ***y***_Λ_*k*__ and corresponding forward operator **E**_Λ_*k*__, and 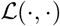 is a loss function. This is supplemented with a self-validation loss on Γ, which is calculated at each epoch *j* from the current network weights specified by ***θ***^(j)^ as follows:

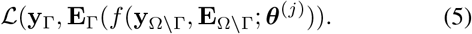

For single dataset training, the loss in Eq. 4 keeps decreasing. The training is stopped once the self-validation loss in Eq. 5 starts increasing to avoid overfitting. A schematic of the ZS-SSDU is depicted in Fig. 1.

**Fig. 1:**
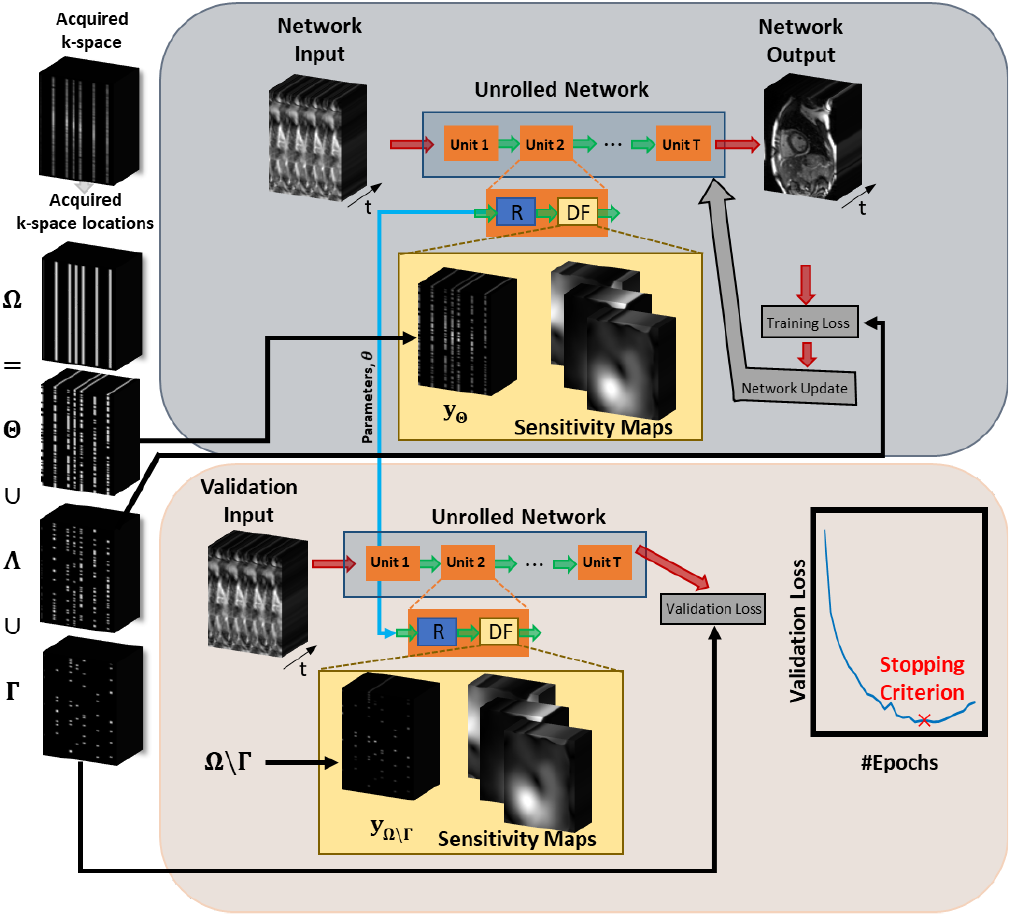
A schematic of zero-shot PG-DL reconstruction. Acquired k-space locations (Ω) are split into three disjoint sets Θ, Λ and Γ. Θ is used in data-fidelity (DF) units of the unrolled network, Λ is used to define k-space self-supervised loss, and Γ is used for k-space self-validation loss to determine a stopping criterion for subject-specific learning. PG-DL network was unrolled for 10 iterations and a 3D ResNet was used as regularizer (R), with all 50 time-frames as input and 128 channels in hidden layers.

## III. Experiments and Implementation Details

### A. Imaging Experiments

Imaging was performed at 1.5T, first at lower resolution with mild acceleration to investigate performance in a retrospective acceleration setting, and then at high resolution with high acceleration to establish translational feasibility.

#### 1) Retrospectively Accelerated Datasets

These were acquired on 11 subjects using a bssFP sequence at accel-eration R = 4 with a Cartesian TGRAPPA sampling pattern [4]. Relevant imaging parameters: FOV=360×270mm^2^, resolution=2.25×2.93mm^2^, and slice-thickness=8mm (11 slices). These datasets were further retrospectively undersampled to R = 7.2 by sampling every 8^th^ line, while keeping the closest *k_y_* line to central k-space for each time-frame.

#### 2) Prospectively Accelerated High-Resolution Datasets

These were acquired on 2 subjects using a Cartesian bssFP sequence with R = 8 acceleration using TGRAPPA sampling, along with corresponding breath-hold cine MRI. Relevant imaging parameters: slice-thickness=8mm (11 slices); realtime: FOV=360×340mm^2^, resolution=1.5× 1.5mm^2^, temporal resolution=44ms; ECG-gated: FOV=360×360mm^2^, resolution=1.4× 1.4mm^2^, temporal resolution=42ms.

### B. Implementation Details

Subject-specific regularization across time-frames was performed using Zs-ssDU [15] using all cardiac cycles. A 3D PG-DL network was unrolled for 10 iterations with a 3D ResNet used for the regularizer where 50 time-frames in its inputs and 128 channels in its hidden layers. Training was performed with Adam optimizer with a learning rate of 4 · 10^-4^ and a mixed normalized *l*_1_ – *l*_2_ loss in k-space [12]. R adjacent frames were used to generate calibration data in a sliding-window manner [4], [5], on which coil sensitivity maps were generated using EsPIRiT. 20% of Ω was uniformly randomly selected for Γ [15] and *K* = 100 was used for disjoint training pairs.

Comparisons were made to TGRAPPA [4], locally-low-rank (LLR) regularized reconstruction [16] and database-trained SSDU [12]. TGRAPPA was implemented using 5 × 4 kernels on calibration data generated from R adjacent frames [4]. For LLR regularized reconstruction, 8 × 8 patches were used with a thresholding value of 0.1 times the *l*_∞_ norm of the zero-filled image. Additionally, for the lower resolution datasets, database training with multi-mask SSDU [22] was used for comparisons. This was trained on 7 subjects using all available slices and 50 time-frames shifted by 20 frames with a total of 365 *k_x_*-*k_y_*-*t* k-spaces over 100 epochs. It used the same network architecture as in ZS-SSDU, and testing was performed on 2 different subjects not used in training. Database training could not be performed for the high-resolution datasets due to the small number of training samples. For lower resolution data, TGRAPPA at standard R = 4 was used as baseline images for comparisons, and PSNR and SSIM were calculated with respect to these images. These were assessed using paired t-test with *P* < .05 considered significant.

## IV. Results

### A. Retrospectively Accelerated Datasets

Fig. 2a shows reconstructed time-frames for R = 7.2 accelerated real-time dataset. TGRAPPA reconstruction at R = 4 acquisition is depicted on the leftmost column for baseline image quality. At R = 7.2, TGRAPPA shows residual artifacts and noise amplification, whereas LLR-regularized reconstruction shows spatio-temporal blurring. Database-trained SSDU shows substantial residual aliasing artifacts due to varying spatio-temporal correlations among subjects in the database. The proposed database-free ZS-SSDU improves all methods, closely matching the baseline R = 4 image. Fig. 2b depicts the difference images to baseline image at R = 4, where proposed methods only show noise-like differences. Temporal signal intensity profiles are shown in Fig. 2c, where database-trained SSDU and TGRAPPA show visible aliasing, while LLR-regularized reconstruction shows substantial temporal blurring. The proposed method closely matches with the baseline profile.

**Fig. 2:**
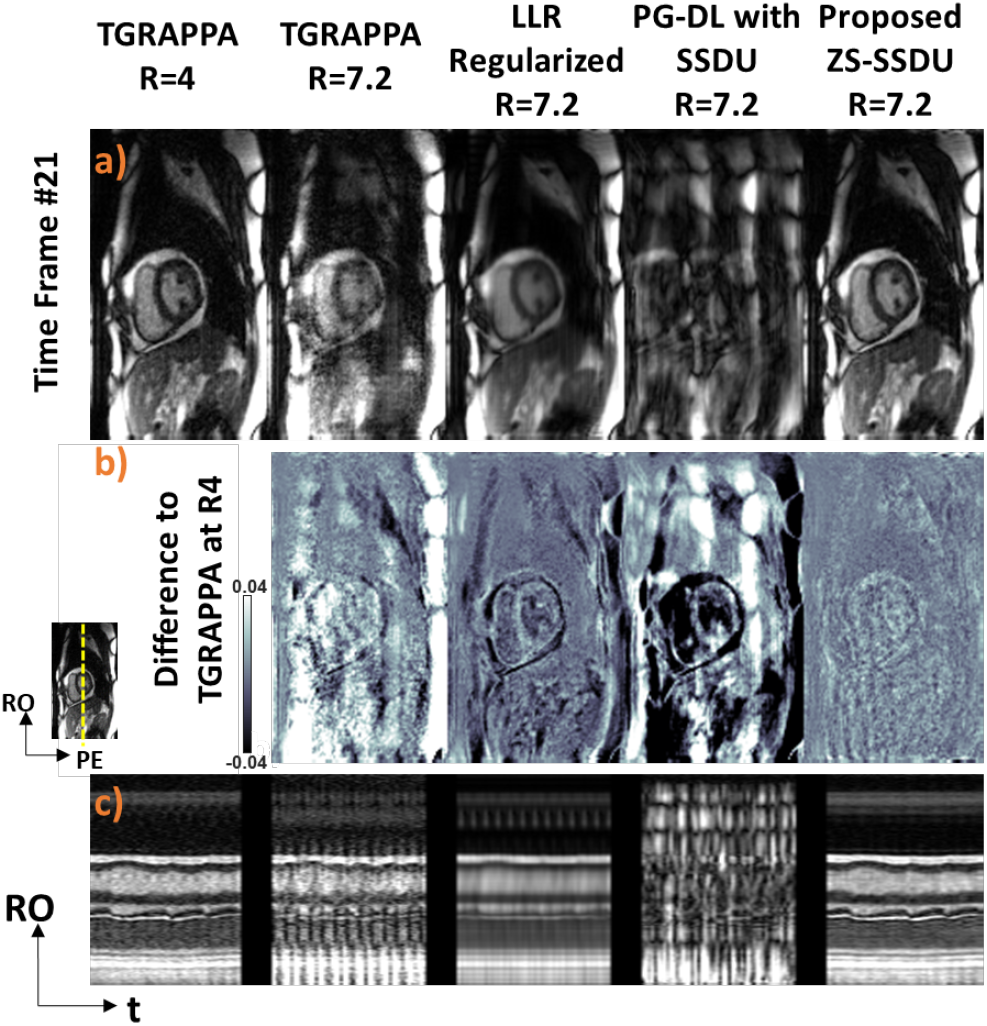
Representative real-time cine MRI results with retrospective acceleration, along with TGRAPP at acquired R = 4 for visual baseline. **(a)** SSDU with database training suffers from severe aliasing artifacts, TGRAPPA shows residual aliasing and noise amplification, whereas LLR regularization exhibits blurring and signal void in the anterolateral segment. Proposed ZS-SSDU at R = 7.2 substantially improves upon all methods, closely matching acquisition acceleration R = 4 data. **(b)** Difference images, with R = 4 TGRAPPA, align with these observations, where our proposed ZS-SSDU only shows noise-like artifacts, while all other methods show structural differences. **(c)** Temporal signal intensity profiles across the blood–myocardium boundary (yellow dashed line) show that proposed ZS-SSDU has similar temporal profiles compared to R = 4 data, with similar cardiac and respiratory motion profiles.

Quantitative PSNR and SSIM metrics across all slices and cardiac cycles for two subjects are in concordance with the previous visual assessments: Database-trained SSDU has the lowest PSNR and SSIM (21.58 ± 3.32 (dB), 54.49 ± 13.80 %), followed by TGRAPPA (22.56 ± 1.75, 73.50 ± 4.06 %) and LLR-regularized reconstruction (28.14 ± 1.29, 81.44 ± 2.83 %). The proposed method outperforms all methods (all pairwise t-tests: P < 10^-4^) with substantial gains in PSNR and SSIM (33.71 ± 1.94, 89.76 ± 2.07 %).

### B. Prospectively Accelerated High-Resolution Datasets

Fig. 3a shows a representative reconstructed time-frame for a prospectively R = 8 accelerated real-time cine MRI acquisition. ECG-gated breath-hold segmented cine MRI is shown as baseline image quality on the leftmost column. As in Section IV-A, TGRAPPA suffers from aliasing, noise and signal void, and LLR-regularized reconstruction exhibits blurring and signal voids. The proposed ZS-SSDU improves upon all methods and shows excellent image quality compared to breath-hold cine MRI. Fig. 3b shows temporal signal intensity profiles, where our proposed ZS-SSDU for R= 8 free-breathing real-time cine MRI closely matches with baseline profile of breath-hold segmented cine MRI. Note since these came from different acquisitions, exact alignment was not possible. Finally, we note database training of PG-DL reconstruction was not performed due to the small number of training subjects.

**Fig. 3:**
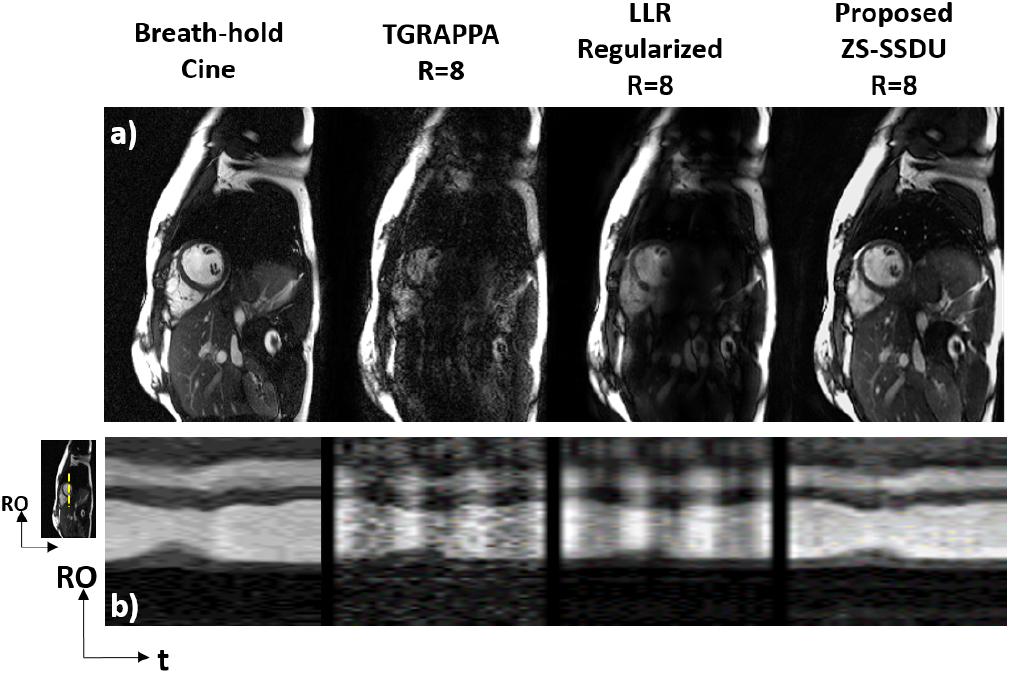
Representative real-time cine MRI results for a prospectively R = 8 accelerated high-resolution dataset. Breath-hold ECG-gated segmented cine MRI from a separete scan is presented on the leftmost column for visual baseline. **(a)** TGRAPPA shows aliasing artifacts and LLR-regularized reconstruction shows signal void and spatial blurring, while proposed ZS-SSDU shows excellent image quality, closely matching to breath-hold segmented cine acquisition. **(b)** Temporal signal intensity profiles show excellent match between the proposed method and breath-hold cine, whereas other methods fail to preserve temporal information.

## V. DISCUSSION

In this study, we proposed a subject-specific zero-shot PG-DL reconstruction with spatio-temporal regularization, which does not require a database to train, for highly-accelerated free-breathing real-time cine cardiac MRI. The main advantage of using ZS-SSDU is addressing robustness and generalizability issues of database-trained models [23] with regards to variations in subject-specific breathing and cardiac motions. The proposed approach improved upon a linear reconstruction, a conventional regularized reconstruction, and database-trained PG-DL reconstruction with better image quality and reduced aliasing.

The database-trained SSDU in this study is trained on 365 k-spaces with shifted time kernels for data augmentation. Even with the data augmentation, the results of database-trained SSDU show residual aliasing indicating database learning of spatiotemporal correlations is difficult. Moreover, the results show that when subject-specific breathing patterns and cardiac motion is presented, database training methods have poor generalizability. Further studies with improved spatio-temporal resolutions in a larger cohort are warranted.

## VI. CONCLUSIONS

The proposed database-free PG-DL reconstruction enabled high-quality spatio-temporal regularization for highly-accelerated real-time Cartesian cine cardiac MRI, learning breathing and cardiac motions reliably without temporal blurring. and improved upon existing techniques, including database-trained PG-DL.

## ACKNOWLEDGMENT

This work was partially supported by NIH R01HL153146, R21EB028369, P41EB027061, NSF CCF-1651825.

## Notes

### Competing Interest Statement

The authors have declared no competing interest.

